# Profiling Unbinding Events to Identify Near-Native RNA-Ligand Poses

**DOI:** 10.1101/2022.02.05.479229

**Authors:** Yichen Liu, Aaron T. Frank

## Abstract

Predicting the structure (or pose) of RNA-ligand complexes is an important problem in RNA structural biology and drug discovery. Although one could use molecular docking procedures to rapidly sample putative poses of RNA-ligand complexes, accurately discriminating the native-like poses from non-native, decoy structures remains a formidable challenge. Here, we started from the assumption that native-like RNA-ligand poses are less likely to dissociate during molecular dynamics simulations, and then we used enhanced simulations to promote ligand for diverse poses of a handful of RNA-ligand complexes. By fitting unbinding profiles obtained from the simulations to a single-exponential, we identified the characteristic decay time (*τ*) as particularly effective at resolving native poses from decoys. Remarkably, a simple regression model trained to predict the simulation-derived parameters directly from structure could also discriminate poses. As such, when molecular dynamics simulations are feasible, one could use them to simulate ligand unbinding to aid in the identification of near-native poses. On the other hand, when speed is more important, one could use a simple structure-based regression model, like the ones described in this study, to analyze and filter RNA-ligand poses.

## INTRODUCTION

Fueled by the approval of the first RNA-targeting small molecule drug,^1^ and the discovery of small bioactive molecules targeting highly structured RNA elements found in microbes,^2–4^ interest in the use of structure-centric methods to identify RNA-targeting small molecules continues to surge. Given a well-validated target, with a clear and demonstrable relationship between its conformational states and biological function,^5^ structure-based methods can, in principle, be used to identify small molecules that bind to and perturb the conformational distribution of that RNA. If one can discover or design such conformationally-specific RNA-binders, they could use them to disrupt RNA function in a therapeutically desirable manner.

One strategy for identifying small molecules that target a specific conformation of RNA is by structure-based virtual screening (SBVS).^6,7^ Structure-based virtual screening involves docking a library of small-molecule ligands to a structure of the target and, on the basis of the docking scores (which are estimates of the binding affinities), sorting them to identify the subset of compounds in the library that are likely to bind to the target. Even though one expects false positives, if one finds true positive hits within the small subset of compounds selected based on docking, the screening is considered successful. However, one of the several assumptions invoked when using SBVS is that the docking calculations can correctly predict the pose (structure) of individual RNA-ligand complexes. This assumption is important because the binding affinity estimations one uses to filter the initial screening library depend on the docked poses; if the docked pose is incorrect, the estimated binding affinity is likely to be incorrect as well. Unfortunately, despite recent progress, the accuracy with which RNA-ligand poses can be predicted lags behind the accuracy currently achievable for protein-ligand complexes. Thus, there is a need to develop new approaches that one can use to complement existing techniques for RNA-ligand pose prediction.

Partly inspired by techniques that leverage molecular dynamics (MD) simulations to aid in protein-ligand^8–11^ and protein-protein^12,13^ pose prediction, we attempted in this study to explore the extent to which one can use information derived from MD simulations to guide pose prediction of RNA-ligand complexes. Here, we leveraged an enhanced sampling method we recently developed, selectively scaled MD (ssMD) simulations,^14^ to promote ligand unbinding during short simulations and from the resulting trajectories generate unbinding profiles of individual RNA-ligand poses. We then assessed the resolving power of parameters derived from these unbinding profiles. Overall, we found that one can use the characteristic decay times (*τ*) of these unbinding profiles to discriminate native from non-native RNA-ligand poses. Furthermore, building off previous studies that show that one can use machine learning to learn from simulations data,^15^ we demonstrated that we could reproduce simulation-derived unbinding profiles directly from the structure of individual poses. Interestingly, the *τ* estimated from structure using the regression model retained the pose resolving power of the *τ* estimated from simulations. We envision that our work should pave the way for using a kinetic framework to assess docked poses of RNA-ligand complexes.

## METHODS

### Preparing Systems for Molecular Dynamics Simulations

Five RNA-ligand systems were studied using selectively scaled MD simulations (ssMD).^14^ The description of the five systems is available in Table S1. For each system, we generated 100 poses using the tool rDock^16^ and the poses generated were ranked based on estimated binding affinities. In order to keep a reasonable computational cost, we reduced the number of poses by clustering. The clustering process was carried out using the Multiscale Modeling Tools for Structural Biology (MMTSB) tool set.^17^ The clustering was based on the cartesian coordinate RMSD between structures, where structures with similar coordinates among heavy atoms were grouped into the same cluster. Different cutoff values were used to keep the number clusters to be less than 20 for each system. For each system, we selected the pose with the best score from each cluster as candidate poses. The structures of the native pose in complex with RNA for ssMD simulations of the 5 RNA-ligand system were obtained from PDB using the PDB ID 2B57, 2XNW, 3NPN, 4XWF, and 5KPY. The structures for the decoy poses were obtained by replacing the coordinates of the native ligand with the decoys’. The input files for the MD simulations were generated using CHARMM-GUI.^18^ To simulate the dynamics of RNA, we used the CHARMM36 nucleic acid force field, and for the ligands, we used the CHARMM General Force Field (CGenFF).^19^ The RNA-ligand systems were solvated in a cubic TIP3P water box with a minimum distance of 10 Å between the solute and the boundary of the water box. Potassium ions and chloride ions were added to neutralize the solvated system.

### Molecular Dynamics Simulations

Each simulation consists of two parts, namely, an equilibration run and a production run. Equilibration simulations were carried out under NVT conditions. SHAKE algorithm was used to constrain hydrogen atoms.^20^ Harmonic restraints were applied to the backbone heavy atoms of RNA with a force constant of 1.0 kcal/mol/Å^2^ and for nucleobase heavy atoms with a force constant of 0.1 kcal/mol/Å^2^. For each equilibration run, the time step was 1 fs and consisted of 2,000,000 steps (or 2 ns). The structures obtained after equilibration were used as the starting structures for the production run. For the production run, hydrogen atoms were constrained using the SHAKE algorithm. Selectively scaled MD (ssMD) simulations have been used to accelerate unbinding events,^14^ where selected energy terms are scaled. For our systems, the Lennard-Jones potential between ligand and water molecules was scaled up to accelerate the dissociation. The assumption is that under stronger ligand-water interactions, the ligand is more likely to dissociate from the binding pocket, thus shortening the simulation time required to observe productive unbinding events. To achieve this, the *ϵ*_*ij*_ term of the Lennard-Jones potential of the ligand and water atoms was scaled up by the scaling factor 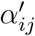.

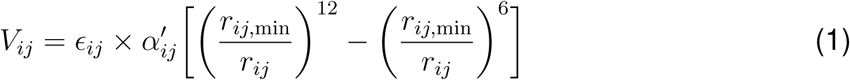

In this work, the scaling factor was set to be 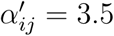. Production simulations were carried out under NPT conditions. For each pose of each of the RNA-ligand systems, we carried out 20 independent 10 ns simulations with a 2 fs time step, so the simulation consists of 5,000,000 steps. Coordinates were written out to a trajectory file every 5000 steps; therefore, each trajectory consisted of 1000 frames. The trajectories obtained from the ssMD simulation were then used for analysis.

### Analyzing Molecular Dynamics Simulations

For each RNA-ligand system, the root mean squared deviations (RMSDs) of the decoy poses relative to the native pose were calculated using PyMOL. Our analysis of the ssMD trajectories focused on the contacts between RNA and ligand. In order to calculate the contacts between RNA and ligand, we prepared a reference contacts file for each pose. The reference contacts were generated using the equilibrated structures with a cutoff value of 3.5 Å. If a pair of atoms in the RNA and the ligand is within 3.5 Å apart, they were considered in contact with each other. We used the contact tool from MoleTools https://github.com/atfrank/MoleTools to calculate the contacts from the ssMD trajectories, where the distances between the atom pairs in the reference contacts were calculated for each frame of the trajectory. Our hypothesis is that the native pose should be more stable compared to decoys, so we generated an unbinding profile for each pose to see if the native poses stay longer than the decoys. For each pose, we calculated the mean number of reference contacts that remain at each frame over the 20 independent trajectories as the unbinding profile. The cutoff value we used to determine whether a contact remains is 3.5 Å. We then fitted the unbinding profile to a decaying exponential function (⟨*M* (*t*)⟩ = *Ae*^(−*t/τ*)^ + *B*), where ⟨*M* (*t*)⟩ is the average number of remaining contacts described as a function of time *t*. The parameters generated from the exponential fitting were then used to assess the binding stability of different poses.

## RESULTS AND DISCUSSION

### Simulation-derived unbinding profiles help resolve native-like poses from decoys

This study was motivated by our interest in developing methods to enhance RNA-ligand pose prediction, that is, methods capable of accurately discriminating native poses from non-native (decoy) poses. If one assumes that the native poses of RNA-ligand complexes correspond to the most stable RNA-ligand conformation, it seems likely that near-native poses will exhibit slower dissociation rates than non-native poses. Based on this assumption, we sought to assess whether native-like poses display distinct unbinding profiles from non-native poses. To address this, we used enhanced molecular dynamics simulations to generate all-atom unbinding trajectories of diverse poses for a set of five benchmark RNA-ligand complexes (Figure 2a). The enhanced sampling method we employed was the selectively scaled MD (ssMD) simulation.^14^ In ssMD, water-ligand interactions are strengthened such that dissociation trajectories can be observed over the course of relatively short simulations without the need for any *a priori* assumptions about the details of the escape paths of the ligand.

**Figure 1:**
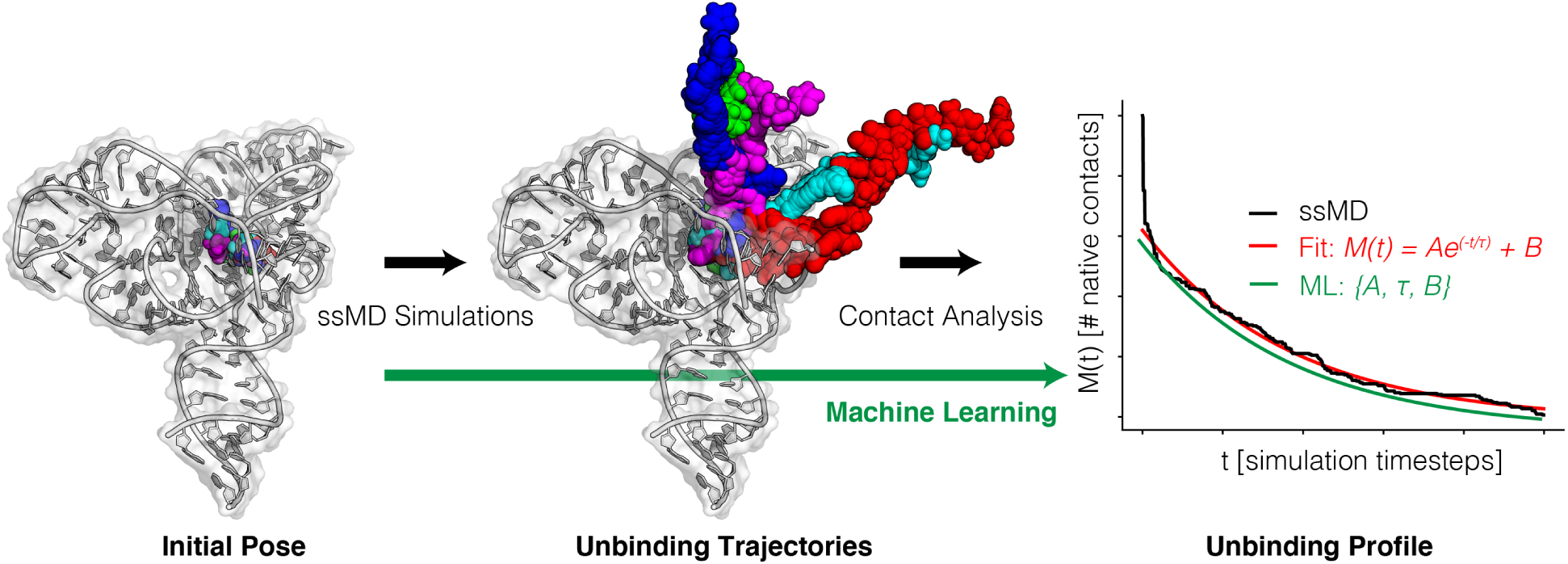
Illustration of the simulation-based framework exploited in this study. For individual poses, we generate a series of unbinding trajectories using the selectively scaled MD (ssMD) technique. We then apply contact analysis to the resulting trajectories and, from them, compute an unbinding profile, defined as the number of native contacts that persist as a function of time. We then fit the unbinding profile to a single exponential function to extract a set of parameters (*A, τ*, and *B*) that compactly describe the unbinding profile. Here we assessed whether we could use any of these parameters to correctly identify near-native poses. In an attempt to bypass costly simulations, we also explored whether we could learn from simulations and predict, directly from the structure of the initial pose, the parameters of the unbinding profile (green arrow).

**Figure 2:**
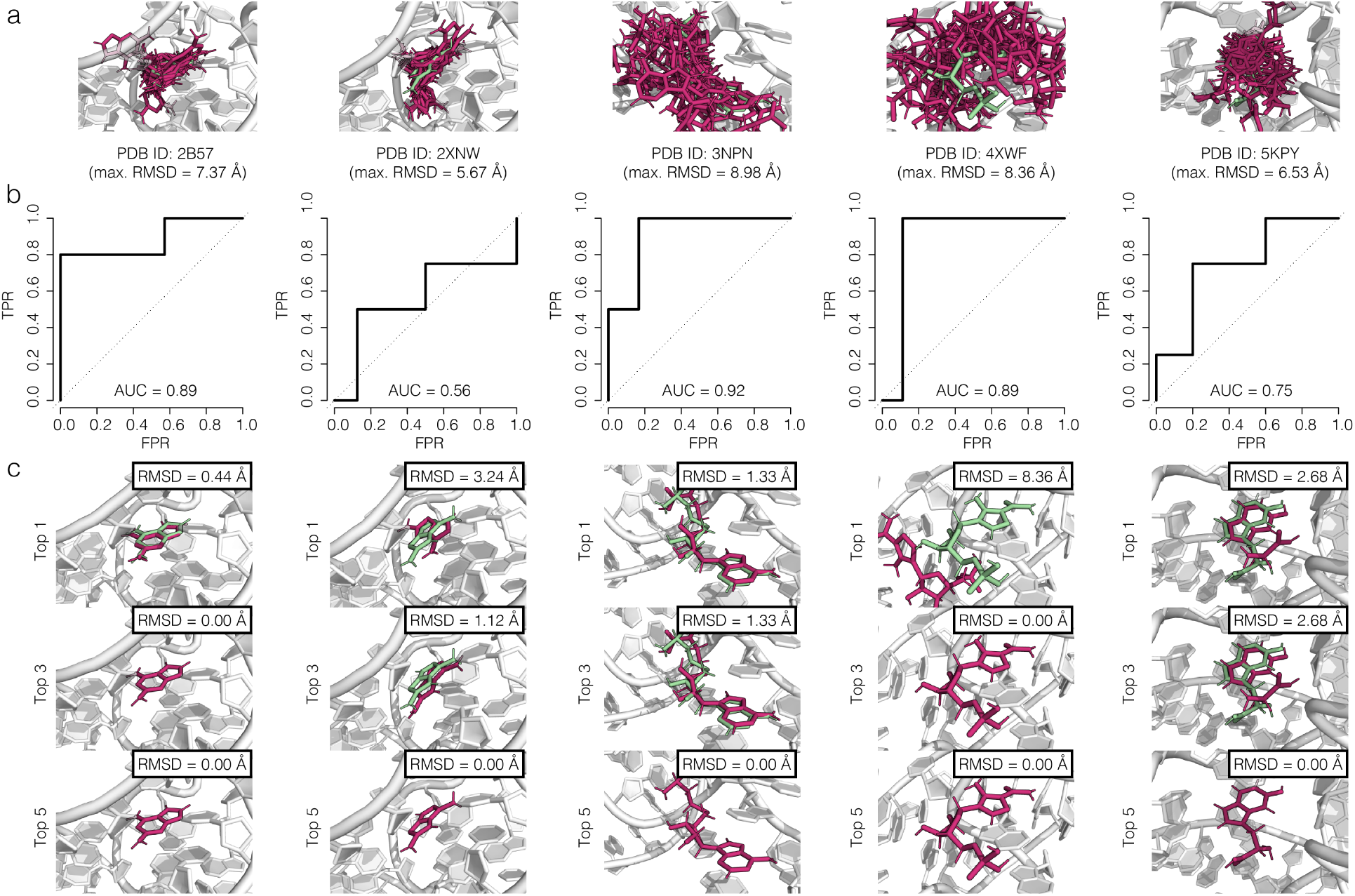
(a) Overlay of poses for each of the five RNA-ligand complexes we examined in this study. (b) Corresponding ROC plots for discriminating native-like poses from non-native (decoy) poses based on the characteristic decay times (*τ*_ssMD_) estimated from ssMD simulations of individual poses. (c) Comparison between the native (green) and best poses (pink) among the top 1, 3, and 5 poses, when we sort poses by *τ*_ssMD_.

Briefly, for each pose in each test case, we generated a set of 20 independent unbinding trajectories using the ssMD method. From the trajectories, we then extracted the time series of all of the (reference) contacts present in the initial pose. Across the 20 replicas, we then computed the mean number of reference contact present at each time step in the save trajectories. The resulting unbinding profile (defined as *M* as function of time (*t*) were then fitted to a decaying, single-exponential function (⟨*M* (*t*)⟩ = *Ae*^(−*t/τ*)^ + *B*).

Across the five test cases, we then carried out receiver-operating-characteristic (ROC) analysis to generate ROC plots when poses are ranked using either *A, B*, or *τ*. In ROC plots, AUC approaching 1 corresponds to instances where a given metric (in the case *A, B*, or *τ* respectively) is able to perfectly separate a set of objects with binary labels, here ‘native-like’ and ‘non-native/decoy’. By contrast, values near 0.5 correspond to what would be expected if labels were randomly assigned to the poses.

In Figure 2b, we show receiver-operating-characteristic (ROC) plots for the five test cases when we used *τ* to rank poses. Among the three fitting parameters, *τ* exhibited the highest resolving power, as judged by the AUC values (Figure S1). Overall the AUC_*τ*_ values ranged between 0.56 and 0.92, with a mean of 0.80 (Table S1). By comparison, AUC_*A*_ and AUC_*B*_, were 0.41 (Figure S1a) and 0.72 (Figure S1c), respectively. Shown in Figure 2c is our comparison between the native poses and the best poses among the top 1, 3, and 5 when they are ranked using *τ*. The RMSD between the native and the pose with the highest *τ* (top 1 prediction) ranged between 0.44 and 8.36 Å, with a mean of 3.21 Å. By comparison the average best RMSD across the five system were 1.03 and 0.00 Å when considering the top 3 and 5 ranked poses (Figure 2c and Table S1). As such, across the five test cases, we could accurately identify near-native RNA-ligand poses by ranking them based on their *τ* derived from ssMD unbinding simulations and identify at most the top five poses.

### Structure-based models can robustly predict simulation-derived unbinding profiles

Though based on a small collection of RNA-ligand complexes, the results reported above indicated that estimating the characteristic time-scale of unbinding during enhanced simulations like ssMD might be a viable strategy for identifying near-native poses. However, placed in the context of docking studies of a large collection of RNA-ligand complexes (as one would typically encounter during virtual screening), carrying out such simulations are impractical; generating the 20 ssMD unbinding trajectories for a single pose require 20 × 9 hours = 180 hours of computing. Thus, to explore whether we could bypass simulations and predict ssMD-derived unbinding profiles directly from structure, we trained a structure-based regression model using the partial least squared (PLS) approach.^21^ Briefly, for each pose of the five RNA in our dataset, we generated molecular (pose) fingerprints^22^ and then paired these features with the exponential fitting parameters from the unbinding profile computed from ssMD simulations that were initialized from that pose. Individual PLS models were then used to predict each parameter (*A, τ*, and *B*) from the pose features.

Shown in Figure 3 are cross-validation root-mean-squared-error of prediction (RMSEP) for each exponential fit parameter as a function of the number of components included in the predictor. Using the optimal number of PLS components, the cross-validation RMSEP for log(*A*), log(*τ*), and log(*C*) were 0.20, 1.12, and 1.44, respectively.

**Figure 3:**
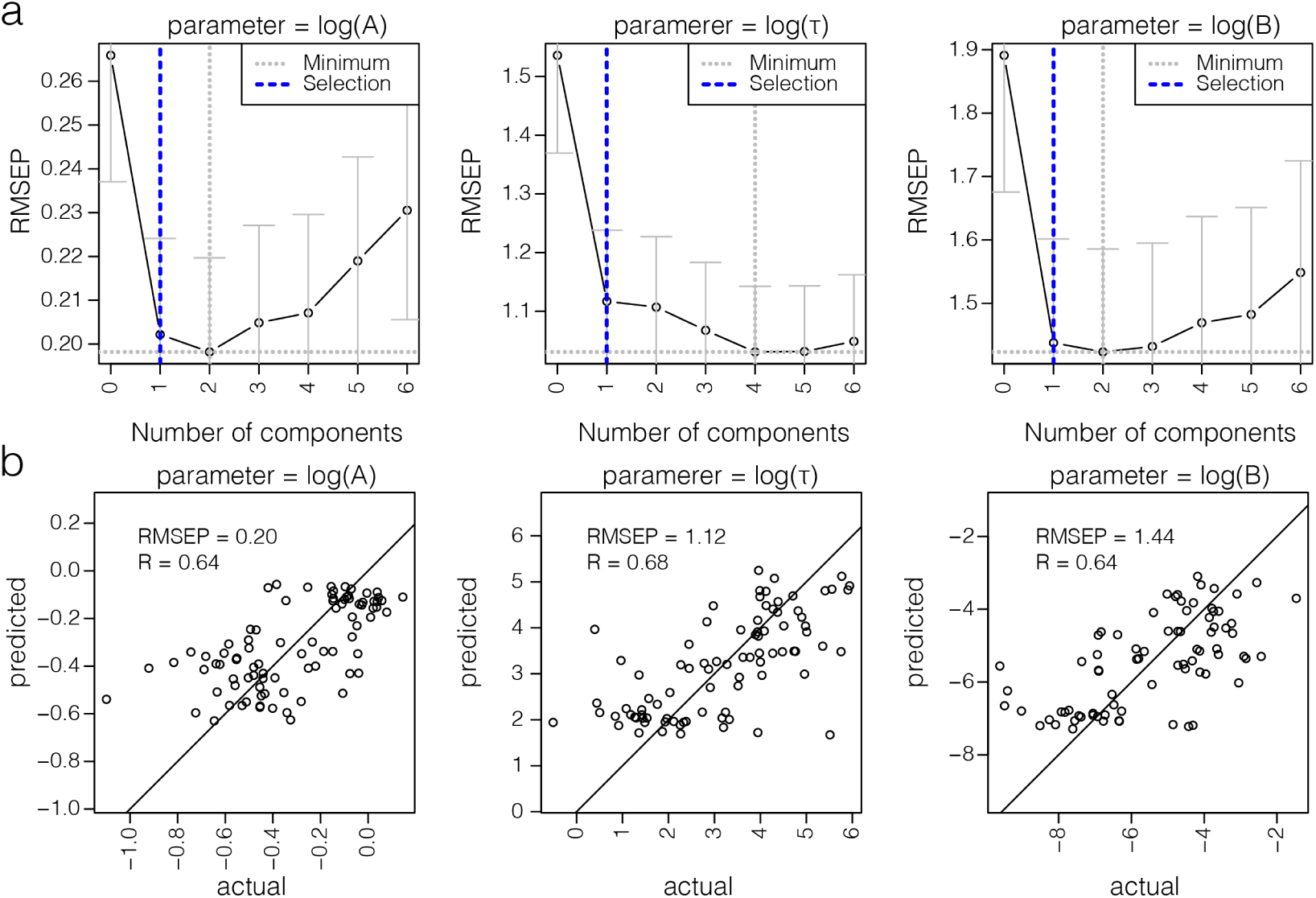
Results we obtained when using partial least squared (PLS) to predict parameters extracted from ssMD unbinding profiles (*A, τ*, and *B*) directly from the structure of individual poses. (a) Cross-validation RMSEP as function of the number of components in the PLS model used to predict log(*A*) (left), log(*τ*) (middle), and log(*C*) (right). Gray dotted lines correspond to the absolute minimum RMSEP observed, and blue dotted lines indicate the optimal selected number of components. We chose the latter using the one-sigma rule as the smallest number of components that exhibited an RMSEP within 1*σ* of the minimum RMSEP. The error bars capture the variance in the estimated RMSEP over the leave-one-out cross-validation experiments. (b) Scatter plots comparing actual (ssMD-derived) log(*A*) (left), log(*τ*) (middle), and log(*C*) (right) with those predicted using optimal number of components. Results correspond to those from leave-one-out cross-validation experiments. And we have annotated the scatter plots with the RMSEP and correlation coefficient (*R*).

Also, the correlation coefficient (*R*) were 0.64, 0.68, and 0.64, respectively. Thus, we could train reasonably robust models to predict the exponential fitting parameters obtained from ssMD data directly from features calculated from the structure of individual RNA-ligand poses.

### Structure-derived unbinding profiles help resolve native-like poses from decoys

Next, we used the PLS model to predict unbinding parameters from a collection of RNA-ligand poses that were distinct from the RNA-ligand pairs used to train the PLS model. Because *τ* estimated from ssMD simulations (*τ*_ssMD_) could be used to resolve poses (Figure 2), we asked whether we also resolve native poses from decoys using *τ* values estimated by our PLS model (*τ*_PLS_).

In Figure 4, we show the ROC plots we obtained when using *τ*_PLS_ to rank poses across ten additional RNA-ligand test cases. We note that these complexes and poses were distinct from those used to train the PLS model. With the exception of PDB ID: 4NYA (AUC = 0.36), the AUC values were all greater than 0.75 and the exhibited mean value was 0.80. Commensurate with the AUC results, the RMSD of the best poses among the top 1, 3, and 5, were typically less than 3.00 Å; across the ten RNA-ligand complexes, the mean top 1, 3, and 5 was 2.86, 2.66, and 2.63 Å, respectively. These results demonstrate that the *τ*_PLS_ we computed directly from structure could indeed be used to resolve poses in the dataset we examined, thus negating the need for computationally demanding simulations.

**Figure 4:**
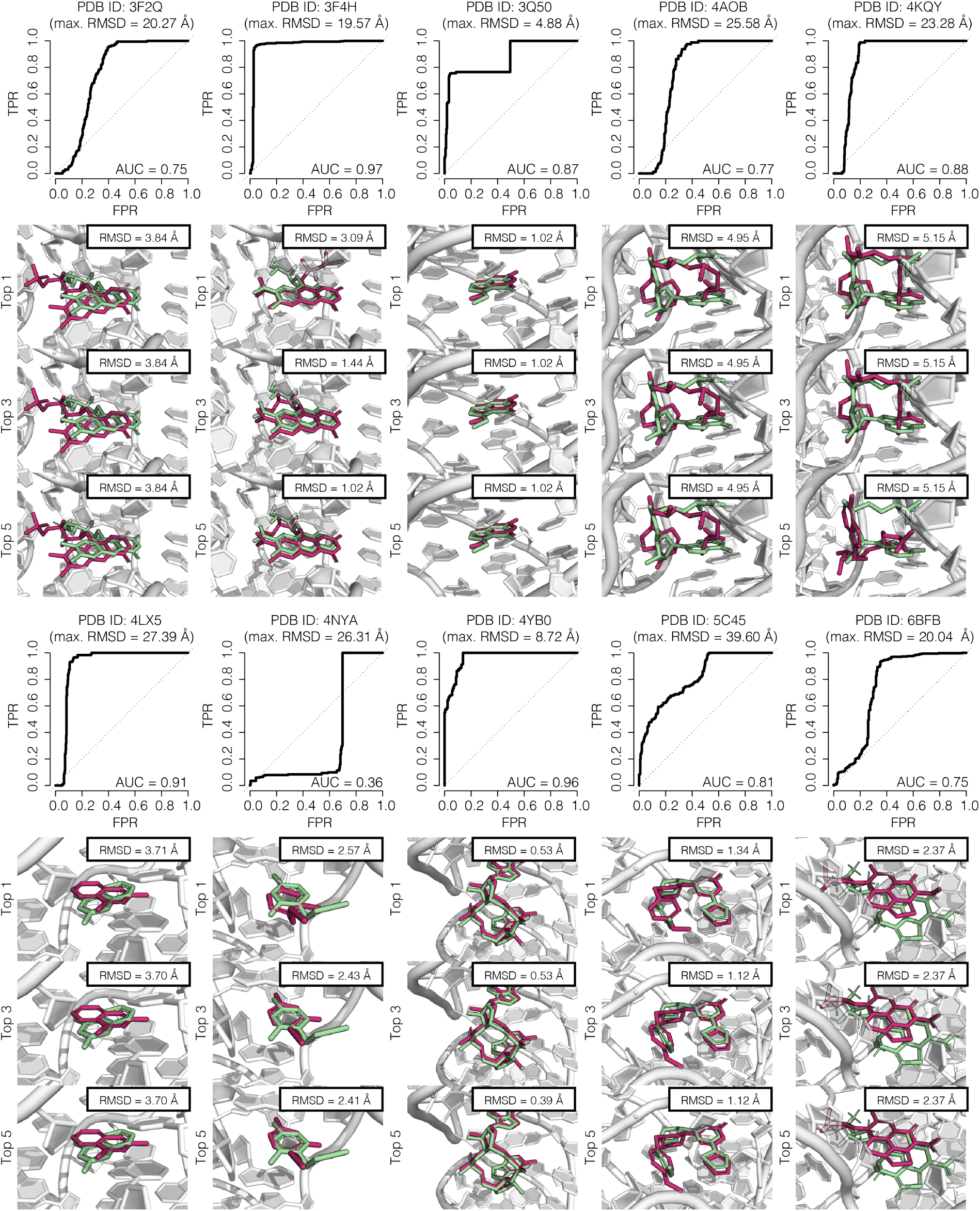
ROC plots for discriminating native-like poses from non-native (decoy) poses based on the characteristic decay times predicted directly from the structure of individual poses using our structure-based PLS model (*τ*_PLS_). Below each ROC plot, we show comparisons between the native (green) and best poses (pink) among the top 1, 3, and 5 poses, when poses are sorted by *τ*_PLS_.

## DISCUSSION AND CONCLUSION

Here we showed that selectively scaled MD simulations can generate unbinding trajectories and that subsequent contact analysis can help identify native-like poses from non-native decoys. This approach is rooted in the comparison of the ligand unbinding profiles of individual poses. When fitted to single-exponential decaying functions, the profiles for near-native poses tend to exhibit longer characteristic decay times (*τ*), and so *τ* could be used to resolve native-like poses from decoys. Building off of these observations, we then showed that we could train a simple PLS regression model to rapidly reproduce the properties of the unbinding profiles (i.e., the parameters of the exponential model) directly from distance-based fingerprints of the initial pose. Though on the basis of our analysis, we found that unbinding parameters estimated from either simulations or a PLS regression model are likely to be useful in resolving RNA-ligand poses, there are a few caveats that must be considered.

First, for our initial ssMD analysis, we carried out multiple independent ssMD simulations for each pose of each RNA-ligand complex. Even with a small database of five RNA-ligand complexes, this amounted to 1720 simulations, each totaling 10 ns and requiring about ∼15,480 hours of computing. Because our initial ssMD-analysis was limited to five RNA-ligand complexes, we cannot rule out that analysis on a larger dataset may yield distinct differing results. However, because we could build a PLS regression model that also exhibited similar resolving power (on a distinct collection of poses) that mirrors that of the ssMD-derived parameters, it gives us some confidence that the trend we observed in our initial ssMD analysis is robust.

Second, though PLS models excel at extracting trends with fewer samples and many features (here model was trained using 86 examples each paired with a 1785-dimensional feature vector), it is likely that increasing the size of the training set might yield a better performing and more generalized model. In this regard, we are currently expanding our database containing ssMD trajectories of unique RNA-ligand poses. We will ultimately use the larger database to update our PLS model.

Third, we restricted our analysis to ligand sensing riboswitches because they represent a subset of RNAs that are highly structured and for which x-ray high-confidence crystal structures are available. Though riboswitches are important antimicrobial drug therapeutic targets in their own right, there is emerging interest in non-riboswitch RNA such as human mRNA and miRNAs as drug targets. However, little to no experimental data is available for such RNA interacting with drug-like molecules. It, therefore, remains an open question whether our PLS model trained using riboswitch data generalizes to such RNAs. Nonetheless, as such data become available, our PLS model could be tested and, if deemed necessary, fine-tuned using data of non-riboswitch RNA-ligand complexes.

Fourth, though *τ* predicted from the PLS model, which was trained on unbinding profiles extracted from ssMD trajectories, exhibited strong resolving power when ranking poses, the absolute *τ*_PLS_ values exhibited a much narrower range than the *τ*_ssMD_ (i.e., the *τ* estimated from the ssMD trajectories). For instance, the mean *τ*_ssMD_ for the native-like poses across training was 153.8 ×10^1^ ps, whereas the mean values predicted from the PLS was 70.5 ×10^1^ ps. By comparison, across the collection of 10 complex systems we used to challenge the PLS model, the values only ranged between 9.0 ×10^1^ ps and 18.0 ×10^1^ ps, with a mean of 11.1 ×10^1^ ps. Therefore, though the relative values *τ*_PLS_ retained resolving power of *τ*_ssMD_, their absolute values may be less meaningful. It is possible that a correction factor could be applied to *τ*_PLS_ to bring them into better correspondence with *τ*_ssMD_; however, given that our interest is really in pose prediction, this is beyond the scope of the current study.

We conclude by noting that despite the limitations/caveats of our approach, the results we have presented strongly indicated that characterizing and quantifying pose dissociation profiles from simulations is a viable strategy for resolving RNA-ligand poses. Moreover, by training regression models that learn from these simulations, it appears that one might be able to bypass these simulations altogether. We envision that such regression models will find utility in the context of docking studies of RNA-ligand complexes. Within the context of such studies, consensus scores can be generated by combining the *τ* that are rapidly predicted using structure-based regression models with scores from other pose prediction strategies.^22–24^ One could then use these consensus scores to identify high-confidence poses that can be used as the input for binding energies estimation and other post-processing virtual screening methods.

## Notes

The code to compute unbinding profiles from structure can be accessed https://github.com/atfrank/RNAPosers/tree/master/ssMD

## Author contributions statement

A.T.F. conceived the project, Y.L. carried out simulations, Y.L. and A.T.F carried out the analysis and wrote and reviewed the manuscript.

## SUPPORTING INFORMATION

**Figure S1:**
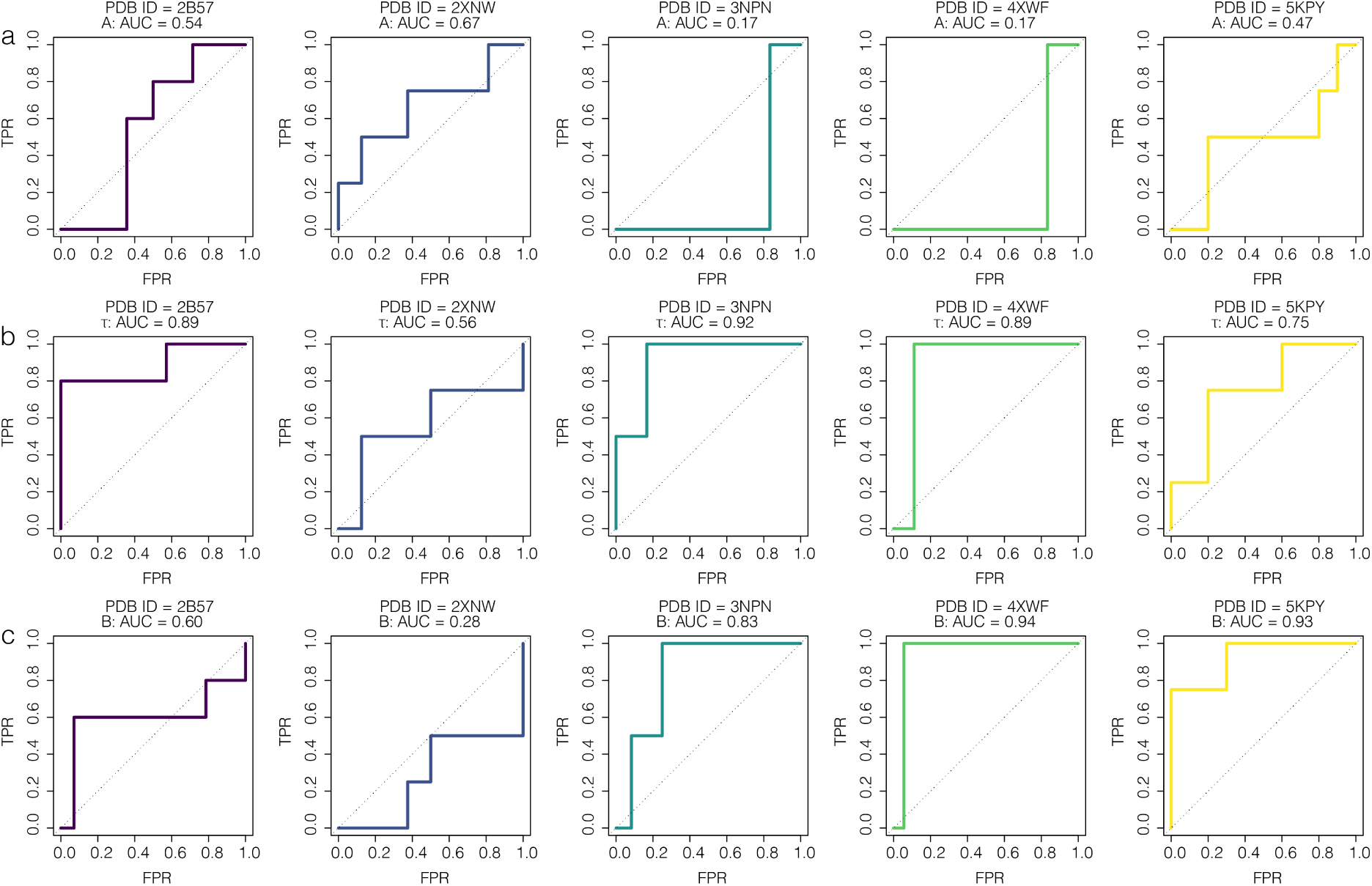
ROC plots when using the ssMD-derived exponential fit parameters (a) *A*, (b) *τ*, and (c) *B* of five RNA-ligand complexes, here identified using their PBD IDs.

**Table S1:** Pose-prediction accuracy metrics when using *τ* estimated from ssMD-derived unbinding profiles to rank RNA-ligand poses. *N* is the number of poses, and the min RMSD listed is the lowest RMSD when considering the top 1/3/5 poses.

| System | Description | $N$ | AUC | min RMSD (Å) |
| --- | --- | --- | --- | --- |
| 2B57 | Purine riboswitch bound to 2,6-Diaminopurine | 19 | 0.89 | 0.44/0.00/0.00 |
| 2XNW | Purine riboswitch bound to triazolo-triazole-diamine | 20 | 0.56 | 3.24/1.12/0.00 |
| 3NPN | SAH riboswitch bound to S-adenosylhomocysteine | 14 | 0.92 | 1.33/1.33/0.00 |
| 4XWF | ZTP riboswitch bound to ZMP | 19 | 0.89 | 8.36/0.00/0.00 |
| 5KPY | 5-hydroxytryptophan aptamer bound to 5-hydroxy-L-tryptophan | 14 | 0.75 | 2.68/2.68/0.00 |
| — | — | 17.2 | 0.80 | 3.21/1.03/0.00 |

**Table S2:** Pose-prediction accuracy metrics when using structure-based *τ*_PLS_ to rank RNA-ligand poses. *N* is the number of poses, and the min RMSD listed is the lowest RMSD when considering the top 1/3/5 poses.

| System | Description | $N$ | AUC | min RMSD (Å) |
| --- | --- | --- | --- | --- |
| 3F2Q | FMN riboswitch bound to FMN | 550 | 0.75 | 3.84/5.50/5.05 |
| 3F4H | FMN riboswitch bound to Roseoflavin | 550 | 0.97 | 3.09/2.00/1.77 |
| 3Q50 | Class I PreQ1 riboswitch to PreQ1 | 500 | 0.87 | 1.02/1.02/1.03 |
| 4AOB | SAM-I riboswitch bound to SAM | 600 | 0.77 | 4.95/8.91/7.49 |
| 4KQY | yitJ S box/SAM-I riboswitch bound to SAM | 600 | 0.88 | 5.15/5.31/5.38 |
| 4LX5 | M6" riboswitch bound to PPDA | 550 | 0.91 | 3.71/3.71/3.71 |
| 4NYA | thiM riboswitch bound 5-(azidomethyl)-2-methylpyrimidin-4-amine | 450 | 0.36 | 2.57/2.58/2.56 |
| 4YB0 | 3',3'-cGAMP riboswitch bound with c-di-GMP | 350 | 0.96 | 0.53/0.54/0.49 |
| 5C45 | FMN riboswitch bound to Ribocil-B | 550 | 0.81 | 1.34/1.22/1.31 |
| 6BFB | FMN riboswitch bound to WG-3 | 550 | 0.75 | 2.37/2.59/2.68 |
| — | — | 525 | 0.80 | 3.21/1.03/0.00 |

